# Characterisation of *in vitro* resistance selection against second-/last-line antibiotics in methicillin-resistant *Staphylococcus aureus*

**DOI:** 10.1101/2024.12.22.630013

**Authors:** Anggia Prasetyoputri, Miranda E. Pitt, Minh Duc Cao, Soumya Ramu, Angela Kavanagh, Alysha G. Elliott, Devika Ganesamoorthy, Ian Monk, Timothy P. Stinear, Matthew A. Cooper, Lachlan J.M. Coin, Mark A. T. Blaskovich

**Author notes:** These authors contributed equally (Anggia Prasetyoputri, Miranda E. Pitt).

## Abstract

**SYNOPSIS:** *Background:* The increasing occurrence of MRSA clinical isolates harbouring reduced susceptibility to mainstay antibiotics has escalated the use of second and last line antibiotics. Hence, it is critical to evaluate the likelihood of MRSA developing clinical resistance to these antibiotics.

*Objectives:* Our study sought to identify the rate in which MRSA develop resistance to vancomycin, daptomycin and linezolid *in vitro* and further determine the mechanisms underpinning resistance.

*Methods:* MRSA was exposed to increasing concentrations of vancomycin, daptomycin, and linezolid for 20 days, with eight replicates for each antibiotic conducted in parallel. The resulting day 20 (D20) isolates were subjected to antimicrobial susceptibility testing, whole genome sequencing, autolysis assays, and growth curves to determine bacterial fitness.

*Results:* Exposure to vancomycin or linezolid for 20 days resulted in a subtle two-fold increase in the MIC, whereas daptomycin exposure yielded daptomycin-nonsusceptible isolates with up to 16-fold MIC increase. The MIC increase was accompanied by variable changes in relative fitness and reduced resistance to autolysis in some isolates. D20 isolates harboured mutations in genes commonly associated with resistance to the respective antibiotics (e.g. *walK* for vancomycin, *mprF* and *rpoB* for daptomycin, *rplC* for linezolid), along with several previously unreported variants. Introduction of key mutations to these identified genes in the parental strain via allelic exchange confirmed their role in the development of resistance.

*Conclusions:* *In vitro* selection against vancomycin, daptomycin, or linezolid resulted in the acquisition of mutations similar to those correlated with clinical resistance, including the associated phenotypic alterations.

## INTRODUCTION

MRSA has retained its ranking as a ‘High’ priority pathogen on the revised 2024 WHO Bacterial Priority Pathogens List^1^ and has been identified as one of the leading causes of global infections^2,3^ and economic burden.^4^ The glycopeptide antibiotic vancomycin has been a mainstay treatment for MRSA-related infections.^5^ Although complete resistance against vancomycin in MRSA is currently rare,^6^ its efficacy is compromised by the increasing incidence of vancomycin-intermediate *S. aureus* (VISA) and heterogeneous VISA (hVISA).^7^ Treatment failures and poor clinical outcomes in glycopeptide-treated patients have been associated with reduced vancomycin susceptibility^6,8,9^ and vancomycin tolerance.^10^ Furthermore, emerging resistance to alternative MRSA antibiotics, such as linezolid^11^ and daptomycin^12^ also adds to the current burden, and highlights the importance for discovery of new MRSA antibiotics.

Clinical resistance in MRSA and its underlying genetic basis has been extensively investigated^6,13–15^ Elucidation of the genomic basis of resistance has been conducted in clinical isolates^16,17^ as well as laboratory-derived strains.^18,19^ Collectively, these have revealed pathways more prone to mutations selected for by clinically-used antibiotics and have been applied to help design new antibiotics that overcome existing resistance mechanisms.

New MRSA antibiotics ideally would have a low propensity to select for clinical resistance. To enable development of such antibiotics, a comprehensive understanding of the nature of resistance mechanisms that might impair their efficacy, and the rate at which that resistance arises, is essential. Availability of *in vitro* resistance selection that could reveal genomic and phenotypic changes similar to those found in clinical settings. This would be a valuable tool for predicting mutations and monitoring the progression of resistance before a candidate antibiotic progressed to the clinic. As an example, laboratory-derived MRSA with reduced susceptibility to daptomycin were generated via resistance selection^20–22^ and certain genomic variants potentially associated with resistance development were identified. Resistance selection could also provide clues to potential cross-resistance between antibiotics having the same cellular target^23^ or give insights into antibiotic resistance mechanisms.^24^

This study aimed to investigate the feasibility of using *in vitro* selection of resistance against vancomycin, daptomycin or linezolid in MRSA to predict mutations similarly identified in clinical isolates. We selected an MRSA strain (ATCC 43300) which is susceptible to vancomycin, daptomycin, and linezolid. ATCC 43300 was serially passaged in the presence of increasing concentrations of each antibiotic and the development of resistance was monitored over 20 days. Genomic and phenotypic alterations were elucidated along with propensities for cross-resistance These investigations are expected to provide further insight into resistance development in MRSA and tools for predicting potential clinical resistance in the context of novel antibiotic discovery.

## MATERIALS & METHODS

### Bacterial strains and growth conditions

MRSA ATCC 43300 isolates were sourced from the American Type Culture Collection (ATCC; Manassas, USA). ATCC 43300 (housed at the University of Queensland - MIC: vancomycin = 1mg/L; daptomycin = 0.5mg/L; linezolid = 2mg/L) was used for *in vitro* resistance selection, subsequent phenotypic assays and genome sequencing. Another ATCC 43300 (University of Melbourne – MIC: vancomycin = 1mg/L; daptomycin = 0.125mg/L) was used in allelic exchange assays. Bacterial isolates were stored in 20% v/v glycerol at -80 °C. ATCC 43300 was grown in cation-adjusted Muller Hinton Broth (Ca-MHB, Oxoid) at 37°C, 220rpm, unless otherwise indicated.

### Resistance selection

*In vitro* resistance selection was conducted^25^for vancomycin, daptomycin, or linezolid, with detailed procedure in Supplementary data. Corning® non-binding surface (NBS) 96-well plates (Sigma- Aldrich) were used to minimise binding of antibiotics to test plate.^26^ (Supplementary material). Following 20 days of antibiotic passaging, isolates were further grown for 5 days (Day 25 (D25)) without antibiotic to assess stability of resistance.

### Antimicrobial susceptibility testing

The MIC was determined by the broth microdilution (BMD) method according to CLSI guidelines^27^ using Ca-MHB (Supplementary data). Susceptibility of all D20 isolates were assessed against their respective selecting antibiotic after the resistance selection experiment was completed, as well as against all antibiotics (vancomycin, daptomycin and linezolid) to assess potential cross-resistance. MIC breakpoints were determined using standards from EUCAST version 14.0,^28^ where resistance was defined as MIC ˃ 2 mg/L for vancomycin, MIC ˃ 1 mg/L for daptomycin, and MIC ˃ 4 mg/L for linezolid.

### DNA extractions and library preparation

Aliquots (10 µL) of glycerol stocks of initial day (day 0/D0) and day 5, 10, 15 and 20 isolates were grown overnight in Ca-MHB supplemented with antibiotics (up to half MIC recorded) (Table S1). Cells from 4 mL of culture were centrifuged (14,000 rpm, 2 mins) and DNA extracted using DNeasy Blood and Tissue Kit (QIAGEN) according to manufacturer’s instructions. Following quantification with Qubit®3.0 (Thermo Fisher Scientific), a library preparation with 1 ng DNA input was conducted using Nextera XT Kit (Illumina). Library preparation of the D0 isolate was performed with the SQK- LSK109 kit (Oxford Nanopore Technologies, ONT) and sequenced on an R9 flow cell. DNA fragmentation was determined with a TapeStation 4200 (Agilent).

### Sequencing and analysis

All DNA libraries were run on Illumina HiSeq2500 with 150 bp paired-end reads and ≥ 100X coverage. The reference D0 isolate was sequenced on a MinION (ONT), base-called using Guppy 2.3.7 and a complete hybrid assembly (Illumina, ONT reads) generated via Unicycler v0.3.7.^29^ Illumina reads were trimmed using Trimmomatic v0.27,^30^ assembled using SPAdes v3.10.1^31^ and annotated using Prokka v1.12.^32^ Trimmed reads were mapped to D0 using BWA-MEM^33^ with default settings. GATK Unified Genotyper^34^ was used to call single-nucleotide polymorphisms (SNPs) and small insertion and deletions (indels) from high-quality reads and impact of non-synonymous variants was annotated using snpEff v4.1,^35^ followed by further quality filtering with SnpSift.^36^

### Autolysis

Selected D20 isolates with increased vancomycin MIC were subjected to Triton X-induced autolysi.^37^ Briefly, overnight cultures were prepared in Brain Heart Infusion (BHI) broth (Oxoid) by inoculating 10µL of glycerol stock into 4mL BHI at 37°C, 200 rpm for 16-20 hours. Subcultures were prepared in 50mL BHI (1:40) and mid log-phase cultures (OD600 = 0.4-0.6) were pelleted (13,000 rpm, 15 mins, 4°C). Cells were washed twice with ice-cold sterile water before resuspension in freshly prepared 0.05 M Tris-HCl (pH 7.2) containing 0.05% (v/v) Triton X-100 (Sigma Chemical Co., St. Louis, Mo.) and incubated at 37 °C, 200 rpm. Absorbance was monitored every 30 min for 5 hours. Results are presented as percentage of OD600 decrease after 5 hours relative to the initial OD600. Controls include D0, MRSA VISA (NRS1; Mu50) ATCC 700699 and MSSA ATCC 29213.

### Bacterial fitness

Relative bacterial fitness was determined by generating growth curves to obtain the average doubling time (DT) following a previous method,^38^ with some modifications (Supplementary data).

### Allelic exchange

To validate if mutations confer resistance, selected genes from vancomycin- and daptomycin D20 isolates were incorporated into WT MRSA ATCC 43300 via allelic exchange as previously described^39^ (Supplementary data, Table S2). All WT and resulting mutant strains underwent antibiotic susceptibility testing as described above.

### Data availability

Nucleotide sequences including genome assemblies (.fasta) and base-called (.fastq) data for both Illumina and ONT are deposited under NCBI BioProject: PRJNA986267 (www.ncbi.nlm.nih.gov/bioproject/986267).

## RESULTS

### Acquisition and retention of *in vitro* resistance differed between antibiotic treatments

The susceptibility profiles of D20 isolates varied greatly between vancomycin, daptomycin, and linezolid following resistance selection (Figure 1a-c). Few vancomycin isolates surpassed the clinical breakpoint (Figure 1a), whereas all of the daptomycin isolates reached the clinical breakpoint of 1 mg/L (Figure 1b). While vancomycin and linezolid isolates experienced less than four-fold increases in MIC after 20 days, daptomycin isolates acquired up to 16-fold increases (Figure 1d). Five days of passaging without antibiotics to discern resistance stability was conducted (Figure 1a-c). Over half of the vancomycin D20 resistant isolates regained susceptibility and all daptomycin isolates retained resistance. Whilst all linezolid D20 isolates were identified as susceptible, re-culturing glycerol stocks revealed three (LZD-2, LZD-3, LZD-4) exhibited resistance (8 mg/L) which was stable for the five antibiotic-free passages.

**Figure 1.**
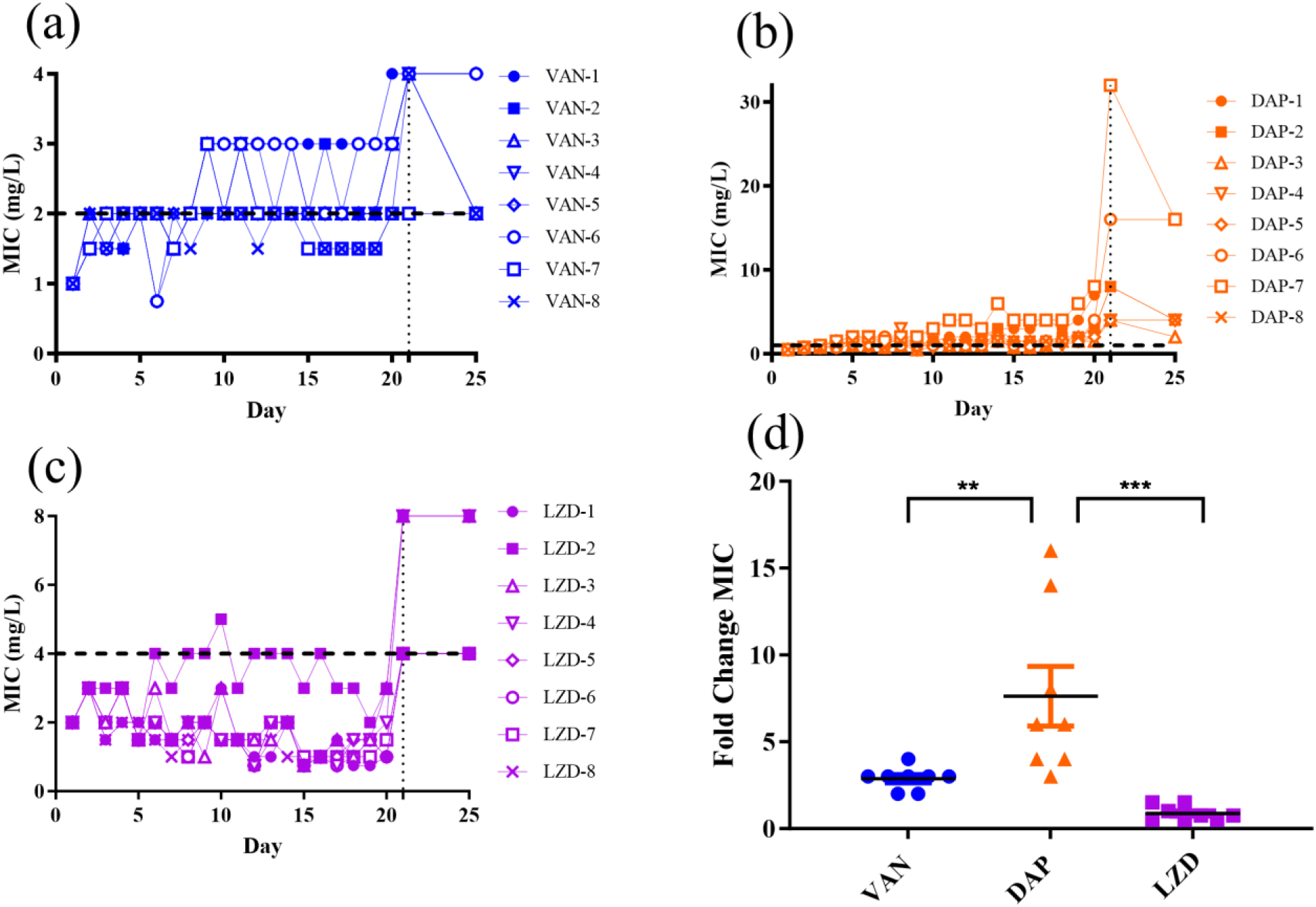
ATCC 43300 acquisition and stability of resistance towards vancomycin, daptomycin, and linezolid. Average MIC values were plotted for (a) vancomycin (VAN)- (b) daptomycin (DAP)- and (c) linezolid (LZD)-treated isolates. Horizontal dotted line represents clinical breakpoints for respective antibiotics based on EUCAST v14.0. Vertical line at D21 includes re-cultured D20 glycerol stocks and subsequent passaging for 5 days with no antibiotics. D25 value indicates highest MIC detected for replicates (n=4). (d) Fold change in MIC values of day 20 isolates compared to day 0 (mean±SD; n=8). Statistical significance was determined by one-way ANOVA with Tukey’s multiple comparisons test (*p* < 0.05 was considered as significant; ** *p* = 0.0080; *** *p* = 0.0003).

### Genomic alterations reflect pathways associated with the antibiotics’ target

Vancomycin isolates exhibited chromosomal mutations in genes associated with cell wall synthesis and metabolism, such as *walK, atl_3* and *korB* (Table 1). Genes involved in DNA repair (*recX*), protein synthesis (*pheS*), and virulence regulation (*tlyC* and *rny*) were also affected. Different mutations occurred across the eight replicates, such as amino acid M246R change in *recX* gene shared by VAN-1 and VAN-7 (Table 2), as well as a deletion in bifunctional autolysin-encoding *atl_3* gene in VAN-5 and VAN-6. No non-synonymous variants were detected in VAN-8; however, resistance was only briefly present in the D20 re-culture and quickly reverted to susceptible (Figure 1a). The frequency these mutations appeared over the time course were monitored (Table S3). The majority of mutations appeared as early as day 5 and established well before D20. Interestingly, the mutations affecting the protein synthesis pathway (*rny* and *pheS*) appeared at a later time and subpopulations having no mutations still existed by D20.

**Table 1.** Non-synonymous variants detected in D20 isolates and the resulting phenotype.

| Strain<sup>a</sup> | Gene name<sup>b</sup> | Protein name | Nucleotide change | Protein change | Phenotype<sup>c</sup><br>(MIC, <u>fold increase from D0</u>) |
| --- | --- | --- | --- | --- | --- |
| VAN-1 | *recX* | Regulatory protein RecX | T737G | Met246Arg | R (4, <u>2</u>) |
|  | *tlyC* | Hemolysin C | C391T | Pro131Ser |  |
|  | *Rny* | Ribonuclease Y | G896A | Arg299Lys |  |
| VAN-2 | *korB* | 2-oxoglutarate oxidoreductase subunit KorB | A386G | Gln129Arg | R (4, <u>2</u>) |
| VAN-3 | **walK** | Sensor protein kinase WalK | C1095A | Asp365Glu | R (4, <u>2</u>) |
| VAN-4 | *pheS* | Phenylalanine—tRNA ligase alpha subunit | C17T | Thr6Ile | R (4, <u>2</u>) |
| VAN-5 | *atl_3* | Bifunctional autolysin | 579delA | Glu193fs | R (4, <u>2</u>) |
| VAN-6 | *atl_3* | Bifunctional autolysin | 579delA | Glu193fs | R (4, <u>2</u>) |
| VAN-7 | *recX* | Regulatory protein RecX | T737G | Met246Arg | S (2, -) |
| DAP-2 | *ptsI* | Phosphoenolpyruvate-protein phosphotransferase | C1399T | Arg467Cys | R (8, <u>8</u>) |
| DAP-3 | *epsJ* | Putative glycosyltransferase EpsJ | G527T | Ser176Ile | R (4, <u>4</u>) |
|  | *ltaS* | Lipoteichoic acid synthase | C296T | Thr99Met |  |
|  | *recX* | Regulatory protein RecX | T737G | Met246Arg |  |
| DAP-4 | *epsJ* | Putative glycosyltransferase EpsJ | G527T | Ser176Ile | R (4, <u>4</u>) |
|  | *ltaS* | Lipoteichoic acid synthase | C296T | Thr99Met |  |
|  | *ychF* | Ribosome-binding ATPase YchF | 426delA | Lys142fs |  |
|  | *recX* | Regulatory protein RecX | T737G | Met246Arg |  |
| DAP-5 | *lacC_2* | Tagatose-6-phosphate kinase | 442_450delGCACAAA<br>TT | Ala148_Ile150del | R (4, <u>4</u>) |
| DAP-6 | *rsmG* | Ribosomal RNA small subunit methyltransferase G | G238T | Ala80Ser | R (16, <u>16</u>) |
| DAP-7 | *rsmG* | Ribosomal RNA small subunit methyltransferase G | G238T | Ala80Ser | R (32, <u>32</u>) |
|  | **mprF** | Phosphatidylglycerol lysyltransferase | C1010T | **Ser337Leu** |  |
|  | *lacC_2* | Tagatose-6-phosphate kinase | 365dupA | Asn122fs |  |
| DAP-8 | **rpoB** | DNA-directed RNA polymerase subunit beta | C2146T | Arg716Cys | R (8, <u>8</u>) |
|  | *lacC_2* | Tagatose-6-phosphate kinase | C833T | Ala278Val |  |
|  | *proS* | Proline--tRNA ligase | T662G | Ile221Ser |  |
| LZD-2 | *rplC* | 50S ribosomal protein L3 | **G463C** | **Gly155Arg** | R (8, <u>2</u>) |
| LZD-3 | *rplC* | 50S ribosomal protein L3 | **G463C** | **Gly155Arg** | R (8, <u>2</u>) |
| LZD-4 | *rplC* | 50S ribosomal protein L3 | **G463C** | **Gly155Arg** | R (8, <u>2</u>) |

<sup>a</sup>VAN: Vancomycin, DAP: Daptomycin, LZD: Linezolid, -: replicate number.
<sup>b</sup>High and moderate impact variants having read depth $\geq 50$ as determined by snpEff. Illumina sequencing produced > 100-fold coverage. fs = frameshift mutation; del = deletion, del = deletion.
<sup>c</sup>Resistant (**R**) based on median MIC (mg/L) and breakpoint according to EUCAST. Fold-increase MIC from day 0 is underlined.
Affected gene(s) and/or mutation(s) previously reported in the literature from clinical and/or laboratory-derived isolates is marked in **bold**.

**Table 2.** MIC of mutants generated by allelic exchange.

| Strain<sup>a</sup> | Vancomycin<br>MIC<sup>b</sup><br>(mg/L) | Fold-change<br>MIC<sup>c</sup> | Daptomycin<br>MIC<sup>b</sup><br>(mg/L) | Fold-change<br>MIC<sup>c</sup> | Parental<br>D20 isolate<br>mutation<br>was detected |
| --- | --- | --- | --- | --- | --- |
| MRSA ATCC 43300 | 1<sup>S</sup> | - | 0.125<sup>S</sup> | - | - |
| *mprF* (Ser337Leu) | 2<sup>S</sup> | 2 | 2<sup>R</sup> | 16 | DAP-7 |
| *rsmG* (Ala80Ser) | 1<sup>S</sup> | - | 0.25<sup>S</sup> | 2 | DAP-6,<br>DAP-7 |
| *ptsI* (Arg476Cys) | 1<sup>S</sup> | - | 0.25<sup>S</sup> | 2 | DAP-2 |
| *walK* (Asp365Glu) | 2<sup>S</sup> | 2 | 0.25<sup>S</sup> | 2 | VAN-3 |

<sup>a</sup>ATCC 43300 isolate (University of Melbourne) was transformed via allelic exchange (MIC: vancomycin - 1 mg/L, daptomycin - 0.125 mg/L) and mutations are depicted as the gene(s) impacted and the respective amino acid changes in brackets. <sup>b</sup>Susceptibility of transformed isolates and parental ATCC 43300 to vancomycin and daptomycin was determined by MIC in Ca-MHB (n =4), where R = resistant and S = susceptible (EUCAST). <sup>c</sup>Fold-change MIC was calculated based on the MIC of transformed compared to parental ATCC 43300.

Daptomycin D20 isolates exhibited more mutations compared to vancomycin, and a greater diversity of pathways were affected (Table 1). Genomic variants were found in genes having a role in cell wall synthesis (*ltaS* and *epsJ*), membrane phospholipid production (*mprF*), protein synthesis pathway (*ychF*, *rpoB*, *proS* and *rsmG*), as well as glucose (*lacC_2*) and carbohydrate (*ptsI*) metabolism pathways. Similar to vancomycin, *recX* (M246R) was also detected in DAP-3 and DAP-4. All isolates became resistant to daptomycin (4-32 mg/L). Development of daptomycin-non-susceptibility (Daptomycin-NS) was likely brought about by an accumulation of mutations, half of which appeared after day 10 (Table S3). The appearance of certain mutations did correspond with increases in MIC. For example, the *mprF* mutation was initially detected at day 10 in the DAP-7 and the MIC increased significantly from day 10 onward (Figure 1b, Table S3). Similarly, the *rpoB* mutation in the DAP-8 started to appear from D15 and by D20 its MIC increased by two-fold (Figure 1b, Table S3).

Variants detected in linezolid isolates were minimal and only impacted *rplC* (G155R) which affects the 50S subunit ribosomal protein L3. Only 3 isolates harboured this mutation and had a resistant phenotype (MIC: 8 mg/L) after re-culturing D20. This mutation appeared between D10-15 in these isolates (Table S3).

### Potential cross-resistance between vancomycin and daptomycin isolates

Nearly all vancomycin D20 isolates exhibited reduced susceptibility to vancomycin, with the exception of VAN-7 (Figure 2a,b) (Table S4). All vancomycin isolates had up to four-fold increase in daptomycin MICs that resulted in daptomycin-NS phenotypes (Figure 2c), suggesting cross- resistance between vancomycin and daptomycin. Conversely, these D20 isolates exhibited a two-fold MIC decrease against linezolid, with all isolates remaining well below the breakpoint (Figure 2d).

**Figure 2.**
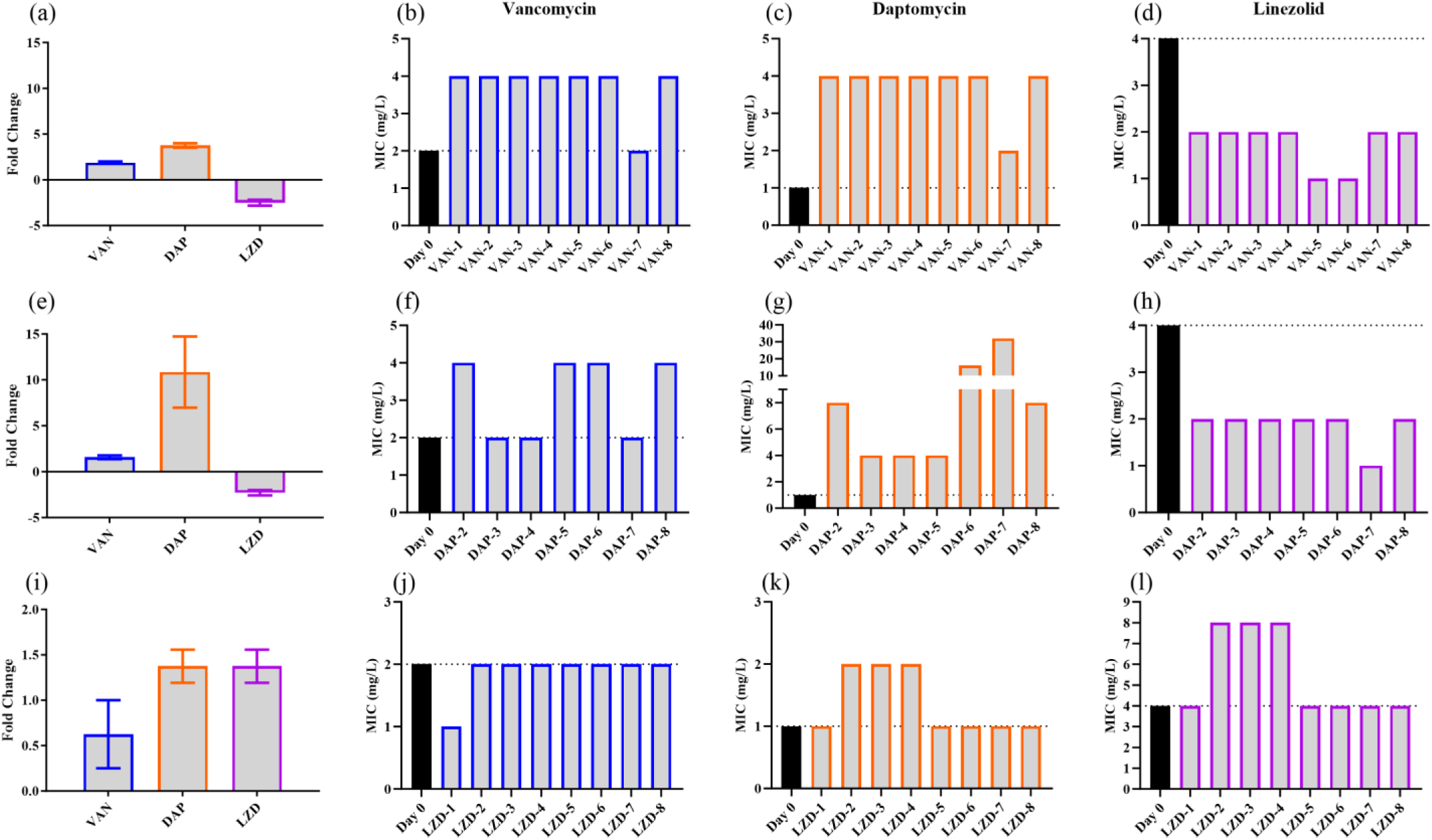
Cross resistance between D20 isolates. (a) Vancomycin D20 isolate fold change for all three antibiotics. MICs of vancomycin D20 isolates against (b) vancomycin, (c) daptomycin, and (d) linezolid. (e) Daptomycin D20 isolate fold change. MICs of daptomycin D20 isolates against (f) vancomycin, (g) daptomycin, and (h) linezolid. (i) Linezolid D20 isolate fold change for all three antibiotics. MICs of linezolid D20 isolates against (j) vancomycin, (k) daptomycin, and (l) linezolid. Fold change represented as average(n=8)±SEM and MICs performed for n ≥ 4. Dotted line indicates breakpoints (EUCAST) and all experiments used re-cultured -80°C D20 glycerol stocks.

Daptomycin D20 isolates exhibited greater than ten-fold increases in MIC against daptomycin (Figure 2e, g). Half of the isolates also had elevated vancomycin MICs over its breakpoint, while the other half exhibited less than two-fold MIC changes (Figure 2e, f). However, the daptomycin D20 isolates remained susceptible to linezolid (Figure 2e, h).

Linezolid D20 isolates demonstrated a slight elevation in MIC against the three antibiotics tested, only by less than two-fold overall (Figure 2i). All D20 isolates remained susceptible to vancomycin (Figure 2j), while three out of eight isolates (LZD-2, LZD-3 and LZD-4) exhibited a daptomycin-NS phenotype, reaching a daptomycin MIC of 2 mg/L (Figure 2k). Although this result suggested potential cross-resistance between linezolid and daptomycin, the linezolid MIC increase was only within a two-fold dilution. The same three isolates also had elevated MICs against linezolid (Figure 2l).

### Variable bacterial fitness costs evident for differing antibiotic exposure

Average doubling time (DT) for each D20 isolate was calculated to assess whether mutations had any impact on the relative bacterial fitness. Vancomycin D20 isolates had a large variability of DT within the biological replicates (Figure 3a) with no significant difference. Daptomycin D20 isolates were found to exhibit DTs longer than that of D0 isolate, though only DAP-2 was significantly different (DT = 69.6±6.1 mins, a 96% increase) (Figure 3b, Table S5). Meanwhile, no significant variation in DTs were found in linezolid isolates (Figure 3c, Table S5).

**Figure 3.**
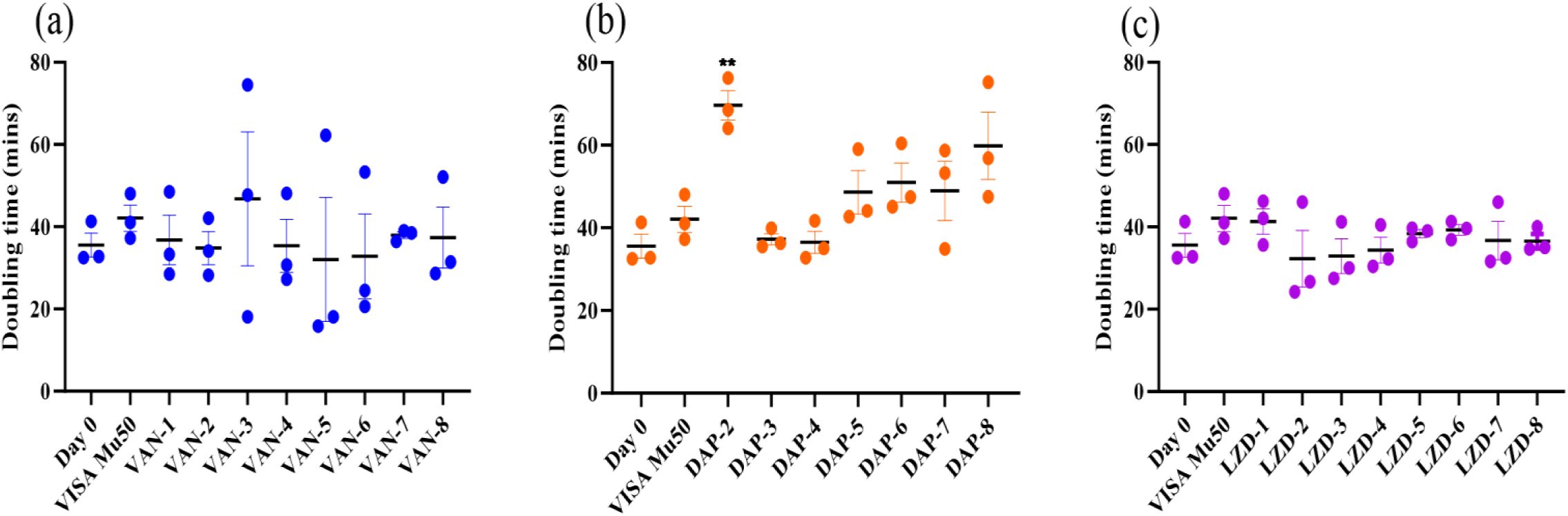
Relative bacterial fitness of D20 isolates. Doubling time (in minutes) of vancomycin (a), daptomycin (b) and linezolid (c) D20 isolates were calculated to measure the relative fitness compared to the initial D0 isolate. Fitness determined using three biological replicates (mean±SD). Significant difference as measured by two-tailed Welch’s t-test (*p*<0.05) between D20 versus D0 is shown with asterisks (\*\**p* = 0.0020).

### Differing degrees of autolysis were observed for vancomycin and daptomycin D20 isolates

D20 isolates were selected based on resistance profile and detected mutations associated with autolysis pathways. The rate of Triton-X-induced autolysis was compared to D0, MSSA and a VISA Mu50 isolate known to exhibit reduced autolysis. VAN-1 and VAN-3 exhibited a slower rate of OD600 decrease compared to D0 with rates similar to the VISA Mu50 (Figure 4a). VAN-8 also had a slower rate of autolysis compared to D0 (50% OD600 decrease after 90 mins) but not as slow as the VISA Mu50 (50% OD600 decrease after 150 mins). In contrast, VAN-5 and VAN-6 had a slightly delayed rate of autolysis compared to D0, though not as slow as the VISA strain, having a 50% OD600 decrease between 60-90 mins. DAP-6 and DAP-7 exhibited similar trends in their resistance to autolysis, having similar rates of OD600 decrease to D0 (Figure 4b). Conversely, DAP-8 showed a higher resistance to autolysis compared to D0, exhibiting similar, if not higher, rate of OD600 decrease to the VISA Mu50. No linezolid isolates were tested for reduced autolysis due to the lack of mutations associated with autolytic activity.

**Figure 4.**
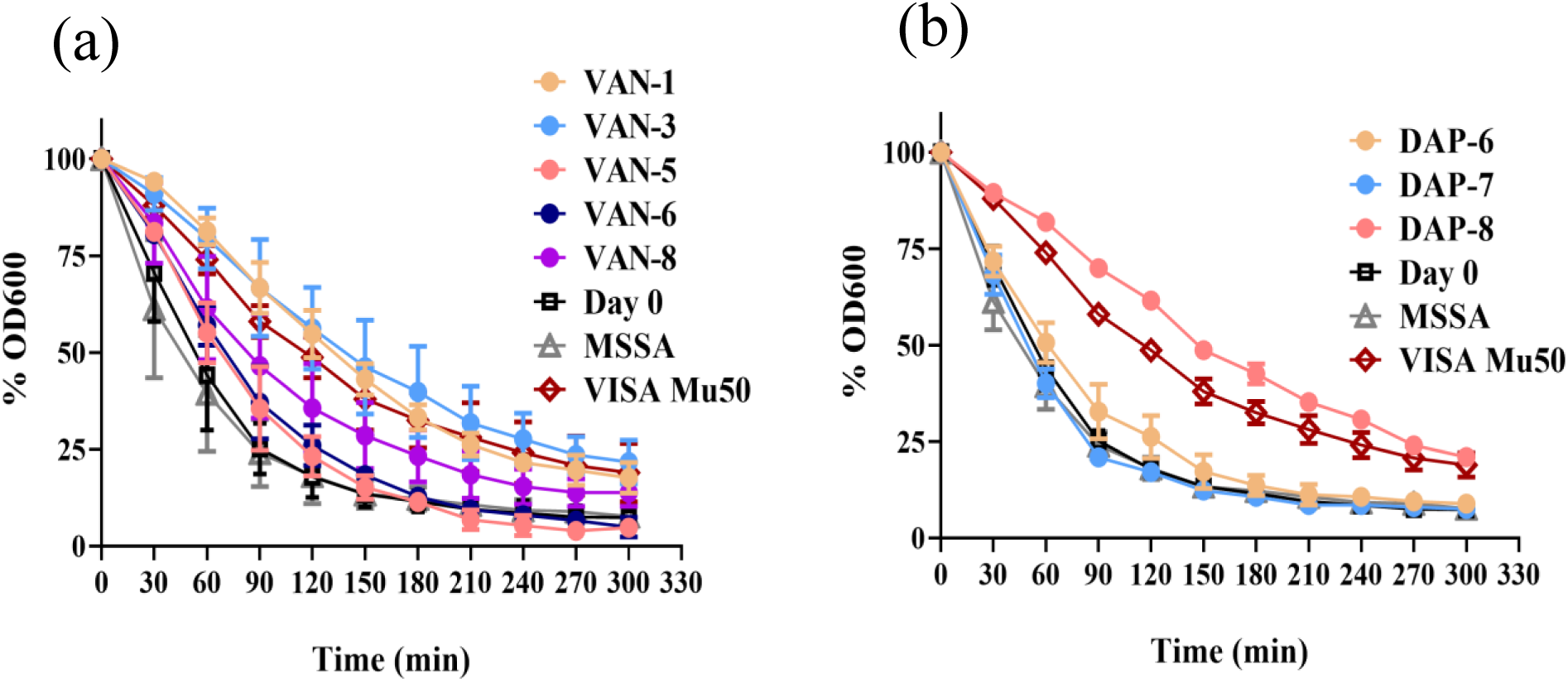
Triton-X-induced autolysis assay on selected vancomycin and daptomycin D20 isolates. Individual plotted percentage of optical density decrease for D20 isolates exhibiting reduced susceptibility. Isolates selected exhibited an elevated MIC as well as mutations in pathways associated with changes in autolysis. (a) Vancomycin isolates VAN-1, VAN-3, VAN-5, VAN-6 and VAN-8 (MIC 4 mg/L); (b) Daptomycin isolates DAP-6, DAP-7, and DAP-8 (MIC 16, 32, and 8 mg/L, respectively). Autolysis measured over five hours. Presented as the percentage decrease in optical density relative to initial D0 (mean±SD). Controls include VISA Mu50 (known tolerance to autolysis) and MSSA (sensitive to autolysis). Data was obtained from three independent experiments.

### Certain variants contribute to the development of resistance

To ascertain whether variants contribute to resistance, we generated mutants of genes harbouring both known and novel mutations via allelic exchange. Four mutants were successfully generated: *walK* (Asp365Glu), *mprF* (Ser337Leu), *rsmG* (Ala80Ser), and *ptsI* (Arg476Cys) (Table 2). Although *rsmG* and *ptsI* have not been reported to be associated with drug resistance in MRSA, *rsmG* (Ala80Ser) was the only change detected in DAP-6 (MIC: 16 mg/L) and present in DAP-7 (MIC: 32 mg/L) (Table 1). The *ptsI* (Arg476Cys) variant was found in DAP-2 exhibiting severely reduced growth rate (Figure 3). The generation of *rpoB* (Gly716Cys) and *rplC* (Gly155Arg) mutants were unsuccessful. As the linezolid mutation in *rplC* was known to cause resistance, this was excluded.

The *walK* (Asp365Glu) mutant had a two-fold increase in vancomycin MIC, which is consistent to our observations in VAN-3 (Table 2). The two-fold increase in daptomycin MIC suggested potential cross-resistance between vancomycin and daptomycin, despite it being less than observed in VAN-3 (four-fold increase in daptomycin MIC compared to D0) (Table 2). Similarly, the *mprF* (Ser337Leu) mutant exhibited a 16-fold increase in daptomycin MIC and a two-fold increase in vancomycin MIC, suggesting cross-resistance and providing evidence of the role of *mprF* (Ser337Leu) in daptomycin- NS phenotype observed in DAP-6.

*rsmG* (Ala80Ser) and *ptsI* (Arg476Cys) had a two-fold increase in daptomycin MIC compared to the parental MRSA, but had no change in their vancomycin MIC. This is different from the MIC determination of the D20 resistance selection isolates, where DAP-2 and DAP-6 had 8-/16-fold increase in daptomycin MIC compared to D0, respectively. The *rsmG* (Ala80Ser) mutation may partially contribute to daptomycin resistance in DAP-7, but further studies are required to unravel the 16-fold increase in DAP-6. Similarly, it is possible that mutations other than the *ptsI* (Arg476Cys) also contributed to the daptomycin-NS phenotype in DAP-2.

## DISCUSSION

*In vitro* resistance selection assays provide key insights into the mechanisms bacteria can employ to develop resistance and can mimic genomic changes detected in a clinical setting. This was evident in our study which exposed MRSA to 20 days of vancomycin, daptomycin or linezolid. Subsequent phenotypes associated with pathways impacted such as fitness and autolysis correlated with clinical isolates.

The WalKR two-component regulator, crucial for cell wall synthesis and metabolism, is strongly associated with vancomycin resistance.^40,41^ Our study also confirmed (VAN-3, *walK* (Asp365Glu), MIC 4 mg/L) and validated this with allelic exchange. The Asp365Glu mutant has yet to be reported and impacts the PAS domain. This variant is within one residue of a lab-derived VISA strain harbouring a *walK* mutation (His364Arg) exhibiting decreased autolytic activity and longer doubling times.^42^

Daptomycin clinical resistance can be associated with genes *mprF* and *rpoB*^43–48^ and *in vitro* assays^20,22,49^, consistent with our findings. *mprF* plays a role in lysinylation of cell membrane phosphatidylglycerol, generating lysyl-PG and its translocation to the outer cell membrane. Mutations impacting the *mprF* gene result in a gain-in-function phenotype that includes increased cell membrane charge, leading to repulsion of the calcium-daptomycin complex.^50^ Interestingly, this particular Ser337Leu mutation in DAP-7 has been reported in several studies associated with the Daptomycin- NS phenotype in clinical and laboratory-derived strains.^43,45,49,51–63^ DAP-7 was found to have the highest increase in daptomycin MIC (32 mg/L), but remained susceptible to vancomycin (MIC: 2 mg/L) as well as a longer doubling time, similar to previous studies.^45,64^ DAP-8 harboured the *rpoB* (Arg716Cys) variant and resulted in reduced susceptibility to both daptomycin (MIC: 8 mg/L) and vancomycin (MIC: 4 mg/L), suggesting cross-resistance. Although the exact mutation has not been reported, the concurrent reduced susceptibility to daptomycin and vancomycin has been reported in other *rpoB* perturbations.^44,65^ *rpoB* mutations can exhibit a VISA phenotype,^66^ which was evident in our study as DAP-8 had an autolysis trend similar to VISA Mu50.

Mutations in *rplC* have been detected in clinical linezolid-resistant isolates^67,68^ Linezolid selection yielded a G463C mutation in *rplC* encoding the 50S ribosomal protein L3. This exact amino acid change has previously been found in an *in vitro* assay using MSSA.^69^ This mutation is located within the peptidyl transferase center (PTC) of the 50S ribosomal subunit, and based on structural studies, can interfere with a conserved part of the PTC and cause reduced binding affinity.^69^ Resistance against linezolid has been largely attributed to mutations in the domain V of 23S rRNA,^12,70^ but mutations in ribosomal protein L3 encoded by *rplC* (as well as ribosomal protein L4 encoded by *rplD*), are also known to be associated with resistance.^70^

Doubling times for all D20 isolates revealed DAP-2 exhibiting a severely impaired growth, in which an Arg476Cys mutation was identified in *ptsI*. *ptsI* is part of the phosphotransferase system (PTS), and mutations in the PTS system have been implicated in heightened daptomycin resistance in *Enterococcus faecium*.^71^ Mutation in *ptsI* is also associated with fosfomycin resistance in *E. coli* that confers a fitness cost.^72^ Further experimentation is needed to confirm whether this *ptsI* mutation is responsible for impaired fitness. Our allelic exchange assay revealed a two-fold increase in daptomycin MIC, suggesting a role in MRSA resistance.

Cross-resistance between vancomycin and daptomycin have often been documented with genes, such as *walK* and *mprF*, which confer both VISA and daptomycin-NS phenotypes.^49,73–75^ For example, mutations affecting *walK* have been shown to cause cross-resistance,^76^ similar to our observations with VAN-3 with daptomycin MIC of 4 mg/L. This is consistent with reports on daptomycin-NS accompanied by reduced susceptibility to vancomycin in the clinic,^77–79^ and daptomycin-NS phenotypes emerging from vancomycin treatment.^79^ This study found that 50% of vancomycin D20 isolates became daptomycin-NS. Similarly, some of daptomycin D20 had elevated vancomycin MIC, as exemplified by DAP-6 harbouring *mprF* (Ser337Leu) causing a two-fold elevation in vancomycin MIC. The concurrent evolution of resistance to both vancomycin and daptomycin may be due to common pathway/s or mechanism/s.^80^ All vancomycin D20 isolates became daptomycin-NS, but daptomycin isolates exhibited variable vancomycin susceptibility profiles. Hence, cross-resistance is likely dependent on the genes and their mutations. Notably, the DAP-7 S337L *mprF* mutation that remained vancomycin susceptible. Ruzin *et al*. demonstrated that *mprF* impairment could lead to increased vancomycin susceptibility.^81^ Additionally, vancomycin reduced susceptibility could be independent to *mprF* alterations.^64^ Yet, other *mprF* mutations can confer reduced susceptibility to both vancomycin and daptomycin.^73^ Alterations in other genes such as *rpoB* could also yield a similar outcome.^65^ Importantly, there could also be implications of the *mprF*-mediated cross-resistance, such as cross-resistance with lipoglycopeptide dalbavancin.^75^

Exposure to sub-inhibitory cell-wall targeting antibiotic concentrations could lead to the development of a VISA phenotype,^82^ which includes autolysis tolerance. The VAN-3 isolate showed reduced autolytic activity similar to a VISA Mu50 strain known to have increased autolysis resistance.^66^ This is consistent with previous reports of mutations affecting the WalKR regulon resulting in a VISA phenotype,^6^ typically characterized by thickened cell wall and reduced autolysis.^66, 83^ Suppression of the autolytic system could result from exposure to sub-inhibitory concentrations of cell wall inhibitors, likely to minimise damage to the cell wall.^84^ No significant difference was observed in the VAN-3 isolate doubling time, as noted for other *walK* mutants.^66^

This study has provided evidence that *in vitro* resistance selection emulated both the genomic and phenotypic changes acquired in the clinic. Although we have confirmed the role of certain genomic variants in contributing to resistance, further studies are needed to validate the role of novel mutations not previously reported, and the effects of multiple mutations in one isolate. Furthermore, this study has provided additional insight into cross-resistance between vancomycin and daptomycin mediated by certain mutations. Identifying variants at the genetic level can provide clues to the underlying mechanisms of these shifts in resistance profiles and provide information for alternative antibiotic treatment in the clinic. Additional experiments providing information on fitness, such as competitive growth assays, would be useful to generate more accurate predictions regarding the likelihood of that mutation being maintained within the population.^85^ This study has demonstrated the feasibility of using *in vitro* resistance selection to predict pathways impacted and in turn, potentially forecast the viability of new antibiotics entering the clinic.

## ACKNOWLEDGEMENTS

The authors would like to thank Dr Arnold Bainomugisa for his guidance on the variant detection pipeline and Ali Hinton for her help with optimising the resistance selection experiments.

## TRANSPARENCY DECLARATIONS

M. A. C. currently holds a fractional Professorial Research Fellow appointment at the University of Queensland with his remaining time as CEO of Inflazome Ltd, a company with headquarters in Dublin, Ireland that is developing drugs to address clinical unmet needs in inflammatory disease by targeting the inflammasome. All other authors: none to declare.

## AUTHOR CONTRIBUTIONS

A.P., M.E.P., A.G.E., M.A.C., L.J.M.C. and M.A.T.B. conceived this study. S.R., and A.K. performed the resistance selection experiments. M.E.P. and D.G. performed the library preparation for WGS. A.P., M.E.P., S.R., A.K., D.G., A.E., and I.M. performed all other microbiological assays, including antibiotic susceptibility testing. A.P., M.E.P. and M.D.C. performed the sequencing analysis. A.P. and I.M. performed the allelic exchange. A.P. wrote the original draft. M.A.C., T.S., L.J.M.C. and M.A.T.B. provided supervision, project administration and funding acquisition. All authors reviewed the data, wrote and revised the manuscript, and approved the final version.

## REFERENCES

1. WHO Bacterial Priority Pathogens List, 2024: bacterial pathogens of public health importance to guide research, development and strategies to prevent and control antimicrobial resistance. Geneva: *World Health Organization*; 2024.

2. Mestrovic T, Aguilar GR, Swetschinski LR et al. The burden of bacterial antimicrobial resistance in the WHO European region in 2019: a cross-country systematic analysis. Lancet Public Health 2022; 7:e897–913.

3. Murray CJ, Ikuta KS, Sharara F et al. Global burden of bacterial antimicrobial resistance in 2019: a systematic analysis. Lancet. 2022;399:629–55.

4. Poudel AN, Zhu S, Cooper N et al. The economic burden of antibiotic resistance: A systematic review and meta-analysis. PLoS One. 2023;18:e0285170.

5. Purrello SM, Garau J, Giamarellos E et al. Methicillin-resistant *Staphylococcus aureus* infections: A review of the currently available treatment options. J Glob Antimicrob Resist 2016; 7: 178–86.

6. Howden BP, Davies JK, Johnson PDR et al. Reduced vancomycin susceptibility in *Staphylococcus aureus*, including vancomycin-intermediate and heterogeneous vancomycin- intermediate strains: Resistance mechanisms, laboratory detection, and clinical implications. Clin Microbiol Rev 2010; 23: 99–139.

7. Blaskovich MAT, Hansford KA, Butler MS et al. Developments in glycopeptide antibiotics. ACS Infect Dis 2018; 4: 715–35.

8. Baek JY, Chung DR, Ko KS et al. Genetic alterations responsible for reduced susceptibility to vancomycin in community-associated MRSA strains of ST72. J Antimicrob Chemother 2017; 72: 2454–60.

9. Holmes NE, Johnson PDR, Howden BP. Relationship between vancomycin-resistant *Staphylococcus aureus*, vancomycin-intermediate *S. aureus*, high vancomycin MIC, and outcome in serious *S. aureus* infections. J Clin Microbiol 2012; 50: 2548–52.

10. Britt NS, Patel N, Shireman TI et al. Relationship between vancomycin tolerance and clinical outcomes in *Staphylococcus aureus* bacteraemia. J Antimicrob Chemother 2017; 72: 535–42.

11. AbdAlhafiz AI, Elleboudy NS, Aboshanab KM et al. Phenotypic and genotypic characterization of linezolid resistance and the effect of antibiotic combinations on methicillin-resistant *Staphylococcus aureus* clinical isolates. Ann Clin Microbiol Antimicrob. 2023;22:23.

12. Nannini E, Murray BE, Arias CA. Resistance or decreased susceptibility to glycopeptides, daptomycin, and linezolid in methicillin-resistant *Staphylococcus aureus*. Curr Opin Pharm 2010; 10: 516–21.

13. Stefani S, Campanile F, Santagati M et al. Insights and clinical perspectives of daptomycin resistance in *Staphylococcus aureus*: A review of the available evidence. Int J Antimicrob Agents 2015; 46: 278–89.

14. McCallum N, Berger-Bächi B, Senn MM. Regulation of antibiotic resistance in *Staphylococcus aureus*. Int J Med Microbiol 2010; 300: 118–29.

15. Howden BP, Peleg AY, Stinear TP. The evolution of vancomycin intermediate *Staphylococcus aureus* (VISA) and heterogenous-VISA*. Infect*, Genet Evol 2014; 21: 575–82.

16. Hafer C, Lin Y, Kornblum J et al. Contribution of selected gene mutations to resistance in clinical isolates of vancomycin-intermediate *Staphylococcus aureus*. Antimicrob Agents Chemother 2012; 56: 5845–51.

17. Mwangi MM, Wu SW, Zhou Y et al. Tracking the in vivo evolution of multidrug resistance in *Staphylococcus aureus* by whole-genome sequencing. Proc Natl Acad Sci 2007; 104: 9451–56.

18. Berscheid A, François P, Strittmatter A et al. Generation of a vancomycin-intermediate *Staphylococcus aureus* (VISA) strain by two amino acid exchanges in VraS. J Antimicrob Chemother 2014; 69: 3190–98.

19. Mishra NN, Yang S-J, Sawa A et al. Analysis of cell membrane characteristics of in vitro-selected daptomycin-resistant strains of methicillin-resistant *Staphylococcus aureus*. Antimicrob Agents Chemother 2009; 53: 2312–18.

20. Friedman L, Alder JD, Silverman JA. Genetic changes that correlate with reduced susceptibility to daptomycin in *Staphylococcus aureus*. Antimicrob Agents Chemother 2006; 50: 2137–45.

21. Song Y, Rubio A, Jayaswal RK et al. Additional routes to *Staphylococcus aureus* daptomycin resistance as revealed by comparative genome sequencing, transcriptional profiling, and phenotypic studies. PLoS ONE 2013; 8: e58469.

22. Camargo ILBdC, Neoh H-M, Cui L, et al. Serial daptomycin selection generates daptomycin-nonsusceptible *Staphylococcus aureus* strains with a heterogeneous vancomycin-intermediate phenotype. Antimicrob Agents Chemother 2008; 52: 4289–99.

23. Arhin FF, Seguin DL, Belley A et al. *In vitro* stepwise selection of reduced susceptibility to lipoglycopeptides in enterococci. Diagn Microbiol Infect Dis 2017; 89: 168–71.

24. Pitt ME, Cao MD, Butler MS et al. Octapeptin C4 and polymyxin resistance occur via distinct pathways in an epidemic XDR *Klebsiella pneumoniae* ST258 isolate. J Antimicrob Chemother 2018; 74: 582–93.

25. Blaskovich MAT, Hansford KA, Gong Y et al. Protein-inspired antibiotics active against vancomycin- and daptomycin-resistant bacteria. Nat Commun 2018; 9: 22–22.

26. Kavanagh A, Ramu S, Gong Y et al. Effects of microplate type and broth additives on microdilution MIC susceptibility assays. Antimicrob Agents Chemother 2019; 63: e01760–18.

27. CLSI, Methods for dilution antimicrobial susceptibility tests for bacteria that grow aerobically; 11th ed. CLSI standard M07. *Clinical and Laboratory Standards Institute*: Wayne, PA, 2018, pp 1–92.

28. The European Committee on Antimicrobial Susceptibility Testing, Breakpoint tables for interpretation of MICs and zone diameters. Version 14.0. http://www.eucast.org, 2024, pp 1–115.

29. Wick RR, Judd LM, Gorrie CL et al. Unicycler: Resolving bacterial genome assemblies from short and long sequencing reads. PLoS Comp Biol 2017; 13: e1005595.

30. Bolger AM, Lohse M, Usadel B. Trimmomatic: a flexible trimmer for Illumina sequence data. Bioinformatics 2014; 30.

31. Bankevich A, Nurk S, Antipov D et al. SPAdes: A new genome assembly algorithm and its applications to single-cell sequencing. J Comput Biol 2012; 19: 455–77.

32. Seemann T. Prokka: rapid prokaryotic genome annotation. Bioinformatics 2014 30: 2068–69.

33. Li H. Aligning sequence reads, clone sequences and assembly contigs with BWA-MEM. arXiv 1303.3997v1 [q-bio.GN] 2013.

34. McKenna A, Hanna M, Banks E et al. The Genome Analysis Toolkit: A map reduce framework for analyzing next-generation DNA sequencing data. Genome Res 2010; 20: 1297–303.

35. Cingolani P, Platts A, Wang LL et al. A program for annotating and predicting the effects of single nucleotide polymorphisms, SnpEff: SNPs in the genome of Drosophila melanogaster strain w(1118); iso-2; iso-3. Fly 2012; 6: 80–92.

36. Cingolani P, Patel VM, Coon M et al. Using *Drosophila melanogaster* as a model for genotoxic chemical mutational studies with a new program, SnpSift. Front Genet 2012; 3: 35–35.

37. Cafiso V, Bertuccio T, Spina D et al. Modulating activity of vancomycin and daptomycin on the expression of autolysis cell-wall turnover and membrane charge genes in hVISA and VISA strains. PLoS ONE 2012; 7: e29573.

38. Lam MMC, Seemann T, Tobias NJ et al. Comparative analysis of the complete genome of an epidemic hospital sequence type 203 clone of vancomycin-resistant *Enterococcus faecium*. BMC Genomics 2013; 14: 595.

39. Monk IR, Stinear TP. From cloning to mutant in 5 days: rapid allelic exchange in *Staphylococcus aureus*. Access Microbiology 2021; 3.

40. Howden BP, McEvoy CRE, Allen DL et al. Evolution of multidrug resistance during *Staphylococcus aureus* infection involves mutation of the essential two component regulator WalKR. PLoS Path 2011; 7: e1002359.

41. Peng H, Hu Q, Shang W et al. WalK(S221P), a naturally occurring mutation, confers vancomycin resistance in VISA strain XN108. J Antimicrob Chemother 2017; 72: 1006–13.

42. Baseri N, Najar-Peerayeh S, Bakhshi B. Investigating the effect of an identified mutation within a critical site of PAS domain of WalK protein in a vancomycin-intermediate resistant *Staphylococcus aureus* by computational approaches. BMC Microbiol 2021; 21: 240.

43. Bayer AS, Mishra NN, Chen L et al. Frequency and distribution of single-nucleotide polymorphisms within *mprF* in methicillin-resistant *Staphylococcus aureus* clinical isolates and their role in cross-resistance to daptomycin and host defense antimicrobial peptides. Antimicrob Agents Chemother 2015; 59: 4930–37.

44. Bæk KT, Thøgersen L, Mogenssen RG et al. Stepwise decrease in daptomycin susceptibility in clinical *Staphylococcus aureus* isolates associated with an initial mutation in *rpoB* and a compensatory inactivation of the *clpX* gene. Antimicrob Agents Chemother 2015; 59: 6983–91.

45. Roch M, Gagetti P, Davis J et al. Daptomycin resistance in clinical MRSA strains is associated with a high biological fitness cost. Front Microbiol 2017; 8.

46. Bayer AS, Mishra NN, Cheung AL et al. Dysregulation of *mprF* and *dltABCD* expression among daptomycin-non-susceptible MRSA clinical isolates. J Antimicrob Chemother 2016; 71: 2100–04.

47. Yang S-J, Xiong YQ, Dunman PM et al. Regulation of *mprF* in daptomycin-nonsusceptible *Staphylococcus aureus* strains. Antimicrob Agents Chemother 2009; 53: 2636–37.

48. Bayer AS, Mishra NN, Sakoulas G et al. Heterogeneity of *mprF* sequences in methicillin-resistant *Staphylococcus aureus* clinical isolates: Role in cross-resistance between daptomycin and host defense antimicrobial peptides. Antimicrob Agents Chemother 2014; 58: 7462–67.

49. Berti AD, Baines SL, Howden BP et al. Heterogeneity of genetic pathways toward daptomycin nonsusceptibility in Staphylococcus aureus determined by adjunctive antibiotics. Antimicrob Agents Chemother 2015; 59: 2799–806.

50. Bayer AS, Schneider T, Sahl H-G. Mechanisms of daptomycin resistance in *Staphylococcus aureus*: role of the cell membrane and cell wall. Ann N Y Acad Sci 2013; 1277: 139–58.

51. Peleg AY, Miyakis S, Ward DV et al. Whole genome characterization of the mechanisms of daptomycin resistance in clinical and laboratory derived isolates of *Staphylococcus aureus*. PLoS ONE 2012; 7: e28316.

52. Jiang S, Zhuang H, Zhu F et al. The Role of *mprF* Mutations in Seesaw Effect of Daptomycin-Resistant Methicillin-Resistant *Staphylococcus aureus* Isolates. Antimicrob Agents Chemother 2022; 66: e0129521.

53. Ji S, Jiang S, Wei X et al. In-Host Evolution of Daptomycin Resistance and Heteroresistance in Methicillin-Resistant *Staphylococcus aureus* Strains from Three Endocarditis Patients. The Journal of Infectious Diseases 2020; 221: S243–S52.

54. Mehta S, Cuirolo AX, Plata KB et al. VraSR two-component regulatory system contributes to *mprF*-mediated decreased susceptibility to daptomycin in *in vivo*-selected clinical strains of methicillin-resistant *Staphylococcus aureus*. Antimicrob Agents Chemother 2012; 56: 92–102.

55. Boyle-Vavra S, Jones M, Gourley BL et al. Comparative genome sequencing of an isogenic pair of USA800 clinical methicillin-resistant *Staphylococcus aureus* isolates obtained before and after daptomycin treatment failure. Antimicrob Agents Chemother 2011; 55: 2018–25.

56. Cameron DR, Jiang J-H, Abbott IJ et al. Draft genome sequences of clinical daptomycin-nonsusceptible methicillin-resistant *Staphylococcus aureus* strain APS211 and its daptomycin-susceptible progenitor APS210. Genome Announc 2015; 3: e00568–15.

57. Rubio A, Moore J, Varoglu M et al. LC-MS/MS characterization of phospholipid content in daptomycin-susceptible and -resistant isolates of *Staphylococcus aureus* with mutations in *mprF*. Mol Membr Biol 2012; 29: 1–8.

58. Kang K-M, Mishra NN, Park KT et al. Phenotypic and genotypic correlates of daptomycin-resistant methicillin-susceptible *Staphylococcus aureus* clinical isolates. *Journal of microbiology (Seoul*, Korea) 2017; 55: 153–59.

59. Quinn B, Hussain S, Malik M et al. Daptomycin inoculum effects and mutant prevention concentration with *Staphylococcus aureus*. J Antimicrob Chemother 2007; 60: 1380–83.

60. Patel D, Husain M, Vidaillac C et al. Mechanisms of in-vitro-selected daptomycin-non- susceptibility in *Staphylococcus aureus*. Int J Antimicrob Agents 2011; 38: 442–46.

61. Steed ME, Hall AD, Salimnia H et al. Evaluation of Daptomycin Non-Susceptible *Staphylococcus aureus* for Stability, Population Profiles, *mprF* Mutations, and Daptomycin Activity. Infectious diseases and therapy 2013; 2: 187–200.

62. Ernst CM, Slavetinsky CJ, Kuhn S et al. Gain-of-Function Mutations in the Phospholipid Flippase MprF Confer Specific Daptomycin Resistance. mBio 2018; 9.

63. Sulaiman JE, Lam H. Novel Daptomycin Tolerance and Resistance Mutations in Methicillin-Resistant *Staphylococcus aureus* from Adaptive Laboratory Evolution. mSphere 2021; 6: 10.1128/msphere.00692-21.6

64. Pillai SK, Gold HS, Sakoulas G et al. Daptomycin nonsusceptibility in Staphylococcus aureus with reduced vancomycin susceptibility is independent of alterations in *mprF*. Antimicrob Agents Chemother 2007; 51: 2223–25.

65. Cui L, Isii T, Fukuda M et al. An rpoB mutation confers dual heteroresistance to daptomycin and vancomycin in *Staphylococcus aureus*. Antimicrob Agents Chemother 2010; 54: 5222–33.

66. Hiramatsu K, Kayayama Y, Matsuo M et al. Vancomycin-intermediate resistance in *Staphylococcus aureus*. J Glob Antimicrob Resist 2014; 2: 213–24.

67. Locke JB, Hilgers M, Shaw KJ. Mutations in ribosomal protein L3 are associated with oxazolidinone resistance in staphylococci of clinical origin. Antimicrob Agents Chemother 2009; 53: 5275–78.

68. Endimiani A, Blackford M, Dasenbrook EC et al. Emergence of linezolid-resistant *Staphylococcus aureus* after prolonged treatment of cystic fibrosis patients in Cleveland, Ohio. Antimicrob Agents Chemother 2011; 55: 1684.

69. Locke JB, Hilgers M, Shaw KJ. Novel ribosomal mutations in *Staphylococcus aureus* strains identified through selection with the oxazolidinones linezolid and torezolid (TR-700). Antimicrob Agents Chemother 2009; 53: 5265–74.

70. Stefani S, Bongiorno D, Mongelli G et al. Linezolid resistance in staphylococci. Pharmaceuticals (Basel) 2010; 3: 1988–2006.

71. Humphries RM, Kelesidis T, Tewhey R et al. Genotypic and phenotypic evaluation of the evolution of high-level daptomycin nonsusceptibility in vancomycin-resistant *Enterococcus faecium*. Antimicrob Agents Chemother 2012; 56: 6051–53.

72. Nilsson AI, Berg OG, Aspevall O et al. Biological costs and mechanisms of fosfomycin resistance in *Escherichia coli*. Antimicrob Agents Chemother 2003; 47: 2850–58.

73. Chen F-J, Lauderdale T-L, Lee C-H et al. Effect of a point mutation in *mprF* on susceptibility to daptomycin, vancomycin, and oxacillin in an MRSA clinical strain. Front Microbiol 2018; 9.

74. Thitiananpakorn K, Aiba Y, Tan X-E et al. Association of *mprF* mutations with cross-resistance to daptomycin and vancomycin in methicillin-resistant *Staphylococcus aureus* (MRSA). Sci Rep 2020; 10: 16107.

75. Hines KM, Shen T, Ashford NK et al. Occurrence of cross-resistance and β-lactam seesaw effect in glycopeptide-, lipopeptide- and lipoglycopeptide-resistant MRSA correlates with membrane phosphatidylglycerol levels. J Antimicrob Chemother 2020; 75: 1182–86.

76. Sulaiman Jordy E, Wu L, Lam H. Mutation in the Two-Component System Regulator YycH Leads to Daptomycin Tolerance in Methicillin-Resistant *Staphylococcus aureus* upon Evolution with a Population Bottleneck. Microbiology Spectrum 2022; 10: e01687–22.

77. Cui L, Tominaga E, Neoh H-m, et al. Correlation between reduced daptomycin susceptibility and vancomycin resistance in vancomycin-intermediate *Staphylococcus aureus*. Antimicrob Agents Chemother 2006; 50: 1079–82.

78. Patel JB, Jevitt LA, Hageman J et al. An association between reduced susceptibility to daptomycin and reduced susceptibility to vancomycin *in Staphylococcus aureus*. Clin Infect Dis 2006; 42: 1652–53.

79. Sakoulas G, Alder J, Thauvin-Eliopoulos C et al. Induction of daptomycin heterogeneous susceptibility in *Staphylococcus aureus* by exposure to vancomycin. Antimicrob Agents Chemother 2006; 50: 1581–85.

80. Chen C-J, Huang Y-C, Chiu C-H. Multiple pathways of cross-resistance to glycopeptides and daptomycin in persistent MRSA bacteraemia. J Antimicrob Chemother 2015; 70: 2965–72.

81. Ruzin A, Severin A, Moghazeh SL et al. Inactivation of *mprF* affects vancomycin susceptibility in *Staphylococcus aureus*. Biochimica et Biophysica Acta (BBA) - General Subjects 2003; 1621: 117–21.

82. Roch M, Clair P, Renzoni A et al. Exposure of *Staphylococcus aureus* to subinhibitory concentrations of β-lactam antibiotics induces heterogeneous vancomycin-intermediate *Staphylococcus aureus*. Antimicrob Agents Chemother 2014; 58: 5306–14.

83. McGuinness WA, Malachowa N, DeLeo FR. Vancomycin resistance in Staphylococcus aureus Yale J Biol Med 2017; 90: 269–81.

84. Antignac A, Sieradzki K, Tomasz A. Perturbation of cell wall synthesis suppresses autolysis in *Staphylococcus aureus*: Evidence for coregulation of cell wall synthetic and hydrolytic enzymes. J Bacteriol 2007; 189: 7573.

85. Martínez JL, Baquero F, Andersson DI. Beyond serial passages: new methods for predicting the emergence of resistance to novel antibiotics. Curr Opin Pharm 2011; 11: 439–45.

