## Supplementary Material for "Characterisation of *in vitro* resistance selection against second-/last-line antibiotics in methicillin-resistant *Staphylococcus aureus*"

[#]These authors contributed equally

### SUPPLEMENTAL MATERIAL

#### This PDF file includes:

Materials & Methods

Tables S1-5

References

#### MATERIALS & METHODS

##### Test compounds

Vancomycin was commercially sourced from Sigma-Aldrich, daptomycin from Molekula, and linezolid from Sapphire Bioscience. Stock solutions were prepared at 12.8 g/L in water for vancomycin and daptomycin, and at 2.56 mg/mL for linezolid. Working solutions at 1.28 g/L concentration (20×) for susceptibility assays were freshly prepared in water.

##### *In vitro* resistance selection

A single colony of MRSA ATCC 43300 was selected and grown overnight in Ca-MHB, shaking at 220 rpm. This overnight culture was then grown to log-phase ($OD_{600}$ = 0.4-0.6) and plated onto separate plates with either vancomycin (Sigma-Aldrich), daptomycin (Molekula) or linezolid (Sapphire Bioscience) (n = 8), with 64 mg/L as the highest concentration. The plates were incubated overnight at 37 °C and bacterial growth monitored for each well. Among the series of antibiotic dilutions, the first well that exhibited the highest growth ($OD_{600} \geq 1$) was diluted (1: 1000) and transferred to a new NBS plate with adjusted range of antibiotic concentrations (depending on change in susceptibility). This procedure was repeated for 20 days (resulting isolates referred to as *day 20 isolates*), followed with five additional days of passage with no antibiotic present to assess reversion of resistance without selective pressure (resulting isolates referred to as *day 25 isolates*). As a control, replicates of bacteria only without antibiotics were passaged in parallel (*day 20 drug-free control*) to enable exclusion of non-antibiotic selected genomic variants. Culture samples from each day were preserved in 20% v/v glycerol and stored at -80 °C. MICs of each antibiotic was recorded daily throughout the course of experiment. Fold change in MIC of the day 20 isolates relative to the day 0 were calculated and significance ($p < 0.05$) was determined using Graph-Pad Prism v8.0.2 with one-way ANOVA and Tukey's multiple comparisons test.

**Antimicrobial susceptibility testing**

For MIC determination of daptomycin, $CaCl_2$ was added to Ca-MHB to a final concentration of 50 mg/L. D0 MRSA ATCC 43300, MRSA VISA (NRS1; Mu50) ATCC 700699 and methicillin-susceptible *S. aureus* (MSSA) ATCC 29213 strains were included as controls. For MIC assays with D20 isolates, aliquots of day 20 glycerol stocks were grown for 20-24 h (37 °C, 200 rpm) in Ca-MHB supplemented with antibiotics at concentrations of up to half the MIC determined at day 20 (Table S1) to maintain resistance. For MIC assays with D25 isolates, no antibiotics were added to the overnight cultures. Compounds (20 ×) were serially diluted two-fold across Corning® NBS 96-well plate (Sigma-Aldrich) resulting in 64 mg/L as the highest concentration tested for each antibiotic. A 40-fold dilution of overnight culture was grown in Ca-MHB until cultures reached $OD_{600}$ = 0.5-0.6. This mid-log phase culture was then diluted to $5 \times 10^5$ CFU/mL and 50 μL added to each well of plates containing test compounds. Sterility (no bacteria) and bacterial growth control (no compound) were included in every plate. Plates were covered and incubated at 37 °C for 24 h. MIC was the lowest concentration showing no visible growth. Results were obtained from at least two independent experiments (four replicates).

MIC of the initial D0 isolate was re-determined (total n = 22) alongside MIC determination of all of the D20 isolates after the resistance selection assay was completed. The resulting median MIC was used as the basis for calculating fold change MIC in subsequent assays. Therefore, vancomycin MIC of D0 was determined to be 2 mg/L, whereas its daptomycin MIC was 1 mg/L and its linezolid MIC was 4 mg/L. D20 isolates were subjected again to MIC determination (each $n \geq 4$) after resistance selection was completed to obtain a set of MICs on which the other phenotypic assays are based. These MICs were redetermined using glycerol stocks of the day 20 isolates that had been re-grown. These re-determined D20 MICs sometimes differed from the MICs obtained directly at the end of the initial 20-day resistance selection experiment depicted in Figure 1. However, any differences in the re-determined MICs from the ones obtained at the end of 20-day resistance selection experiment remained within a one-fold dilution. The 'redetermined' MICs of D0 and D20 isolates will be the values used for comparison throughout this paper (Figure 2).

**Relative bacterial fitness**

Relative bacterial fitness was determined by generating growth curves to obtain the average doubling time following a previous method,[1] with some modifications. Glycerol stock aliquots of D20 isolates were grown overnight in Ca-MHB broth for ~20 h (37 °C, 200 rpm). D0 MRSA ATCC 43300, MRSA VISA (NRS1; Mu50) ATCC 700699 and MSSA ATCC 29213 strains were included as controls. $OD_{600}$ of overnight cultures were measured and culture concentrations were adjusted to $10^7$ CFU/mL by diluting in different amounts of Ca-MHB broth based on their respective $O_{600}$ values. Diluted cultures were then plated on a Corning® 96-well clear polystyrene plate (Sigma-Aldrich) (n=4). The plate was incubated at 37 °C (without shaking) for 20 h in a Tecan M1000 microplate reader (Thermo Fisher Scientific), while bacterial growth was simultaneously monitored by $OD_{600}$ measurement every 30 minutes. Sterility (media only) and bacterial growth control (no antibiotics) were also included. Results were obtained from three independent experiments/biological replicates and analysed using GraphPad Prism v8.0.2. The average doubling time of each isolate was calculated from growth curves.[2] Briefly, $OD_{600}$ values of the technical replicates were averaged and plotted on a log scale graph. Time points marking the beginning (time 0) and the end (time 1) of exponential phase for all isolates were noted to determine the generation time. The final doubling time (in mins) was then calculated by averaging the times obtained for each biological replicate using the following equations:

$$\textit{Generation time}\ (\mathbf{Gt}) = \frac{\log10\ (\mathbf{Time\ 1}) - \log10\ (\mathbf{Time\ 0})}{\log10\ (2)}$$

$$\Delta \textit{Time} = \text{Time 1} - \text{Time 0}$$

$$\textit{Doubling time}\ (\mathbf{Dt}) = (\mathbf{Gt} \times \mathbf{\Delta Time}) \times 60$$

Significance in doubling time was assessed by two-tailed Welch's t-test ($p < 0.05$).

**Allelic exchange**

Allelic exchange to generate *S. aureus* with mutations detected in D20 isolates was performed according to a previously described protocol.[3] The gene of interest was amplified using the AF-BR and CF-DR primer pairs (Table S2) and the desired nucleotide change incorporated. The resulting amplicon was ligated into the plasmid pIMAY-Z. Electrocompetent ATCC 43300 (generated as previously described[3]) was transformed in 1 mm electroporation cuvette (Bio-Rad) at room temperature under the following conditions: 21 kV/cm, 100 ohms, 25 µF (GenePulsar, Biorad). Transformed cells were plated on BHI containing chloramphenicol (Chloramphenicol 10µg / Xgal 100 µg/ ml) overnight at 37°C. Transformed isolates underwent antimicrobial susceptibility testing as previously described to determine the extent the mutation was contributing to resistance.

**Table S1. Concentration of antibiotics for supplementing growth medium in cross-resistance studies and DNA extractions**

| Strain[a] | Antibiotic concentration (mg/L)[b] |
|---|---|
| D0 | 0 |
| VAN-1 | 2 |
| VAN-2 | 1.5 |
| VAN-3 | 1.5 |
| VAN-4 | 1.5 |
| VAN-5 | 1.5 |
| VAN-6 | 1.5 |
| VAN-7 | 1 |
| VAN-8 | 1 |
| DAP-2 | 1.5 |
| DAP-3 | 0.75 |
| DAP-4 | 1 |
| DAP-5 | 1 |
| DAP-6 | 2 |
| DAP-7 | 4 |
| DAP-8 | 1.5 |
| LZD-1 | 0.5 |
| LZD-2 | 1.5 |
| LZD-3 | 1.5 |
| LZD-4 | 1 |
| LZD-5 | 0.5 |
| LZD-6 | 0.5 |
| LZD-7 | 0.75 |
| LZD-8 | 0.75 |

[a]D0 = Day 0 initial strain is MRSA ATCC 43300; D20 isolates are vancomycin- (VAN), daptomycin- (DAP) and linezolid- (LNZ) selected isolates, with numbers indicating the replicate number. DAP-1 could not be recovered for DNA extraction and cross-resistance studies, thus, excluded from all experiments.

[b]Antibiotic concentrations were calculated as half of the MIC value determined for that replicate at the end of resistance induction experiment (day 20).

**Table S2. Oligos used for allelic exchange**

| Gene | 5' – 3' sequence | Source |
|---|---|---|
| IM1 | GGTACCCAGCTTTTGTTCCCTTTAGTGAGG | Prior study[3] |
| IM2 | GAGCTCCAATTCGCCCTATAGTGAGTCG | Prior study[3] |
| IM3 | AATACCTGTGACGGAAGATCACTTCG | Prior study[3] |
| IM4 | TACATGTCAAGAATAAACTGCCAAAGC | Prior study[3] |
| walK_AF | CCTCACTAAAGGGAACAAAAGCTGGGTACCAGGTCGAAACGAATGAA GTGGCTAAAAC | This study |
| walK_BR | GACCTCGTGTAACACAGCGATATAACC | This study |
| walK_CF | GGGTTATATCGCTGTGTTACACGAGGTCACTGAACAACAACAAGTTGA ACG | This study |
| walK_DR | CGACTCACTATAGGGCGAATTGGAGCTCCTCCTTATTATTCATCCCAAT CACCGTC | This study |
| walK_screenR | GGTTATATCGCTGTGTTACACGAGGTC | This study |
| mprF_AF | CCTCACTAAAGGGAACAAAAGCTGGGTACCATGAATCAGGAAGTTAAA AACAAAATATTTTC | This study |
| mprF_BR | CAGCAATGATGGTATTTTAGCAATAATATCTTTTTG | This study |
| mprF_CF | ATATTATTGCTAAAATACCATCATTGCTGTTGGCAATTTTAGTATTCTTT ACAAG | This study |
| mprF_DR | CGACTCACTATAGGGCGAATTGGAGCTCACAAATCACTGACATGTTGA AGTTC | This study |
| mprF_screenR | TGTAAAGAATACTAAAATTGCCAACAGC | This study |
| pstI_AF | CCTCACTAAAGGGAACAAAAGCTGGGTACCCAAGGTATCGGCTTATAT AGAACTGAG | This study |
| pstI_BR | ACACGCAGCTAATGTGTATTGAATTAAATCATTTG | This study |

| | | |
|---|---|---|
| pstI_CF | TTTAATTCAATACACATTAGCTGCGTGTCGTATGTCAGAGCGTGTATC | This study |
| pstI_DR | CGACTCACTATAGGGCGAATTGGAGCTCTAGCTATCATAAATTAAATAT<br>CGATTTTTAA | This study |
| pstI_screenR | ATACACGCTCTGACATACGACAC | This study |
| rsmG_AF | CCTCACTAAAGGGAACAAAAGCTGGGTACCCATCAATTGAATGCAGAT<br>GTTGAAGAAC | This study |
| rsmG_BR | TGAACCAGCGCCTACATCACATATACTTATAGGCTGATTA | This study |
| rsmG_CF | TAAGTATATGTGATGTAGGCGCTGGTTCAGGTTTTCCAAGTATTCCGTT<br>AAAAAT | This study |
| rsmG_DR | CGACTCACTATAGGGCGAATTGGAGCTCTAAAGGATTATGCATTATTTT<br>TCAAGTAAAG | This study |
| rsmG_screenR | AACGGAATACTTGGAAAACCTGAA | This study |

**Table S3. Frequency of reads mapping to alternative allele throughout time course for variants detected in D20 isolates**

| Strain | Gene | Nucleotide change | Amino acid change | Day 5 | Day 10 | Day 15 | Day 20 |
|---|---|---|---|---|---|---|---|
| | | | | ALT%[a] | | | |
| VAN-1 | *tlyC* | C391T | Pro131Ser | 97.2 | 100 | 100 | 100 |
| | *rny* | G896A | Arg299Lys | 0 | 0 | 4.4 | 73.2 |
| | *recX* | T737G | Met246Arg | 98.4 | 100 | 100 | 100 |
| VAN-2 | *korB* | A386G | Gln129Arg | 0.7 | 1.5 | 100 | 90.4 |
| VAN-3 | *walK* | C1095A | Asp365Glu | 99.5 | 94.3 | 100 | 98.5 |
| VAN-4 | *pheS* | C17T | Thr6Ile | 0 | 2.4 | 32.2 | 60.9 |
| VAN-5 | *atl_3* | 579delA | Glu193fs | 0 | 100 | 100 | 100 |
| VAN-6 | *atl_3* | 579delA | Glu193fs | 99.2 | 100 | 100 | 100 |
| VAN-7 | *recX* | T737G | Met246Arg | 0 | 0 | 76.9 | 95.5 |
| DAP-2 | *ptsI* | C1399T | Arg467Cys | 0.4 | 11.6 | 100 | 100 |
| DAP-3 | *epsJ* | G527T | Ser176Ile | 0 | 0.5 | 100 | 100 |
| | *ltaS* | C296T | Thr99Met | 45.7 | 94.7 | 100 | 100 |
| | *recX* | T737G | Met246Arg | 100 | 100 | 100 | 100 |
| DAP-4 | *epsJ* | G527T | Ser176Ile | 0 | 0 | 100 | 100 |
| | *ychF* | 426delA | Lys142fs | 0 | 0.4 | 7.5 | 95.4 |
| | *ltaS* | C296T | Thr99Met | 0 | 93.8 | 100 | 100 |
| | *recX* | T737G | Met246Arg | 0 | 100 | 100 | 100 |
| DAP-5 | *lacC_2* | 442_450delGCACAAATT | A148_I150del | 0.5 | 0 | 100 | 100 |
| DAP-6 | *rsmG* | G238T | Ala80Ser | 0 | 0 | 0 | 71.2 |
| DAP-7 | *mprF* | C1010T | Ser337Leu | 0 | 97.1 | 100 | 100 |
| | *rsmG* | G238T | Ala80Ser | 99.6 | 100 | 99.6 | 100 |
| | *lacC_2* | 365dupA | Asn122fs | 0 | 0 | 0 | 80.5 |
| DAP-8 | *rpoB* | C2146T | Arg716Cys | 0 | 0 | 1.2 | 100 |
| | *lacC_2* | C833T | Ala278Val | 0 | 0.9 | 23.6 | 100 |
| | *proS* | C662G | Ile221Ser | 0.8 | 92.2 | 96.2 | 98.8 |
| LZD-2 | *rplC* | G463C | Gly155Arg | 9.2 | 100 | 99.5 | 100 |
| LZD-3 | *rplC* | G463C | Gly155Arg | 0 | 0 | 0.4 | 99.6 |
| LZD-4 | *rplC* | G463C | Gly155Arg | 0 | 0 | 0.4 | 100 |

[a]Percentage of reads mapping to the alternative (mutation) allele. Shading indicates when alternative allele is ≥50%.

**Table S4. Minimum inhibitory concentrations of D20 isolates against vancomycin, daptomycin and linezolid compared to initial D0**

| Strain[a] | MIC[b] (mg/L) | | |
|---|---|---|---|
| | VAN | DAP | LZD |
| D0 | 1 | 0.5 | 2 |
| VAN-1 | 4 | 4 | 2 |
| VAN-2 | 4 | 4 | 2 |
| VAN-3 | 4 | 4 | 2 |
| VAN-4 | 4 | 4 | 2 |
| VAN-5 | 4 | 4 | 1 |
| VAN-6 | 4 | 4 | 1 |
| VAN-7 | 2 | 2 | 2 |
| VAN-8 | 4 | 4 | 2 |
| DAP-2 | 4 | 8 | 2 |
| DAP-3 | 2 | 4 | 2 |
| DAP-4 | 2 | 4 | 2 |
| DAP-5 | 4 | 4 | 2 |
| DAP-6 | 4 | 16 | 2 |
| DAP-7 | 2 | 32 | 1 |
| DAP-8 | 4 | 8 | 2 |
| LZD-1 | 1 | 1 | 4 |
| LZD-2 | 2 | 2 | 8 |
| LZD-3 | 2 | 2 | 8 |
| LZD-4 | 2 | 2 | 8 |
| LZD-5 | 2 | 1 | 4 |
| LZD-6 | 2 | 1 | 4 |
| LZD-7 | 2 | 1 | 4 |
| LZD-8 | 2 | 1 | 4 |

[a]D0 = Day 0 initial strain is MRSA ATCC 43300; day 20 isolates are vancomycin- (VAN), daptomycin- (DAP) and linezolid- (LNZ) treated isolates, with numbers indicating the replicate number. Blue boxes represent MICs of day 20 isolates against the inducing antibiotic. MIC assays of the day 20 isolates were performed independent of the resistance selection experiment. Results presented as median value from at least two independent experiments in duplicate ($n \geq 4$). [b]Resistance (red shading) was defined as MIC > 2 µg/mL for VAN, MIC > 1 µg/mL for DAP, and MIC > 4 µg/mL for LZD according to EUCAST guidelines version 14.0.

**Table S5. Doubling times of D20 isolates as a result of exposure to vancomycin, daptomycin, or linezolid**

| Strain[a] | Doubling time (DT)[b] (mins) | MIC (mg/L) against the selecting antibiotic[c] | %DT increase compared to D0[d] |
|---|---|---|---|
| D0 | 35.5±5.0 | - | - |
| VISA | 42.1±5.5 | - | - |
| VAN-1 | 36.8±10.4 | 4 | 3.7 |
| VAN-2 | 34.8±7.0 | 4 | - |
| VAN-3 | 46.8±28.2 | 4 | 31.8 |
| VAN-4 | 35.3±11.2 | 4 | - |
| VAN-5 | 32.0±26.2 | 4 | - |
| VAN-6 | 32.8±17.8 | 4 | - |
| VAN-7 | 38.0±1.4 | 2 | 7 |
| VAN-8 | 37.4±12.8 | 4 | 5.4 |
| DAP-2 | 69.6±6.1 | 8 | 96.1** |
| DAP-3 | 37.2±2.3 | 4 | 4.8 |
| DAP-4 | 36.5±4.6 | 4 | 2.8 |
| DAP-5 | 48.6±9.0 | 4 | 36.9 |
| DAP-6 | 50.9±8.2 | 16 | 43.4 |
| DAP-7 | 48.9±12.5 | 32 | 37.7 |
| DAP-8 | 59.9±14.1 | 8 | 68.7 |
| LZD-1 | 41.3±5.4 | 4 | 16.3 |
| LZD-2 | 32.3±11.9 | 8 | - |
| LZD-3 | 32.9±7.3 | 8 | - |
| LZD-4 | 34.3±5.4 | 8 | - |
| LZD-5 | 38.4±1.7 | 4 | 8.2 |
| LZD-6 | 39.3±2.2 | 4 | 10.7 |
| LZD-7 | 36.7±8.1 | 4 | 3.4 |
| LZD-8 | 36.6±3.0 | 4 | 3.1 |

[a]D0 = Day 0 initial strain is MRSA ATCC 43300; VISA strain is Mu50 NRS1 MRSA; D20 isolates are vancomycin- (VAN), daptomycin- (DAP) and linezolid- (LNZ) selected isolates, with numbers indicating the replicate number. [b]Mean doubling times from three independent experiments. [c]Median MICs of D20 isolates determined after the resistance selection experiment was completed, from at least two independent experiments in duplicates ($n \geq 4$). [d]%DT increase compared to D0, DTs less than Day 0 DT were indicated with (-). **Significant difference as measured by two-tailed Welch's t-test (with $p<0.05$ considered significant) between the D20 isolates compared to D0 ($p = 0.0020$).

#### REFERENCES


1. Lam MMC, Seemann T, Tobias NJ et al. Comparative analysis of the complete genome of an epidemic hospital sequence type 203 clone of vancomycin-resistant *Enterococcus faecium*. BMC Genomics 2013;14:595.
2. Baines SL, Holt KE, Schultz MB et al. Convergent adaptation in the dominant global hospital clone ST239 of methicillin-resistant *Staphylococcus aureus*. mBio 2015;6:e00080-15.
3. Monk IR, Stinear TP. From cloning to mutant in 5 days: rapid allelic exchange in *Staphylococcus aureus*. Access Microbiology 2021;3: 000193.
